# An Empirical Bayes Approach for the Identification of Long-range Chromosomal Interaction from Hi-C Data

**DOI:** 10.1101/497776

**Authors:** Qi Zhang, Zheng Xu, Yutong Lai

## Abstract

Hi-C experiments have become very popular for studying the 3D genome structure in recent years. Identification of long-range chromosomal interaction, i.e., peak detection, is crucial for Hi-C data analysis. But it remains a challenging task due to the inherent high dimensionality, sparsity and the over-dispersion of the Hi-C count data matrix.

We propose EBHiC, an empirical Bayes approach for peak detection from Hi-C data. The proposed framework provides flexible over-dispersion modeling by explicitly including the “true” interaction intensities as latent variables. To implement the proposed peak identification method (via the empirical Bayes test), we estimate the overall distributions of the observed counts semiparametrically using a smoothed EM algorithm, and the empirical null by discrete curve fitting. We conducted extensive simulations to validate and evaluate the performance of our proposed approach and applied it to real datasets. Our results suggest that EBHiC can better identify peaks than Fit-Hi-C in terms of accuracy, biological interpretability, and the consistency across biological replicates.

## 1 Introduction

The 3D organization of the chromosomes plays an important role in gene regulation, and its identification has been under spotlight in functional genomics. Following the pioneering work on Chromosome Conformation Capture (3C) by Dekker *et al*. (2002), many types of high throughput experiment protocols have been developed, such as Chromosome Conformation Capture Carbon Copy (5C) (Dostie *et al.*, 2006), Hi-C (Lieberman-Aiden *et al.*, 2009) and *in situ* Hi-C (Rao *et al.*, 2014). Among these technologies, Hi-C types of assays allow genome-wide comprehensive investigation of all genomic loci, and have become the most popular approaches for studying chromatin structure.

Hi-C experiments are designed to generate paired-end reads connecting two genomic loci with physical contact in 3D space, and a Hi-C dataset is usually presented as a contact map, a large sparse matrix whose element in the *i*th row and *j*th column represents the number of reads connecting the *i*th and the *j*th genomic loci which are typically genomic bins with the same fixed size. Depending on the resolution of the data, and the scale of the biological questions, there are many ways of modeling the 3D physical interaction between potential regulatory elements. Researchers have made important findings in analyzing Hi-C data, including the inference on the large-scale structure of the genome such as the topologically associated domains (TADs, e.g.,Rieber and Mahony 2017), the A/B compartments (Zheng and Zheng, 2017) and the 3D spatial structure (Park and Lin, 2017), and the identification of the individual genomic loci pairs with physical interactions (peak calling, e.g., Ay *et al*. 2014; Jin *et al*. 2013).

We are interested in peak detection in this paper, as it is immediately relevant to the central question of functional genomics, which is how to understand the regulatory elements in non-coding region. Take genome-wide association studies (GWAS) and expression quantitative trait loci mapping (eQTL) for example. These analysis have led to many important biological discoveries and created tremendous resources for future biomedical studies. However, the majority of these hits fall outside the protein coding regions or the promoter regions, making the the functional interpretation of their roles in gene regulation difficult. Recent studies suggested that this dilemma can be alleviated by taking into account the chromosomal physical contact in 3D space. For example, Smemo *et al*. (2014) utilized 4C assay, and discovered the long-range interactions between an obesity associated intronic variant in gene FTO and the promoter of the homeobox gene IRX3 whose expression is directly associated with the body mass.

Many peak identification algorithms have been developed for Hi-C data. The mainstream strategy in the literature is to assume a parametric model on the off-diagonal elements of the contact matrix, estimate the individual null parameters, and assign a p-value for each loci pair. Duan *et al*. (2010) assumed a binomial model for each bin pair, with the total read depth as the number of trials. Then the null contact probability was calculated using all the loci pairs with the same genomic distance. Ay *et al*. (2014) proposed the peak calling method Fit-Hi-C. It inherited the same Binomial model, but improved the loci-pair grouping, utilized smoothing spline for null probability estimate, and added a refitting step. Mifsud *et al*. (2017) developed another binomial based peak caller whose null probability was based on the product of the relative coverage of the two loci. Over-dispersion is a ubiquitous phenomenon in high-throughput sequencing data, and Hi-C is no exception. Thus it has been incorporated in Hi-C peak detection methods such as Jin *et al*. (2013), where the null models are Negative Binomial distributions with a common *ad hoc* over-dispersion parameter for all loci pairs.

In this paper, we present an Empirical Bayes model for peak detection from HiC data (EBHiC). Our proposed semi-parametric empirical Bayes approach offers principled probability distribution estimates for HiC counts, and provides flexible modeling of over-dispersion by explicitly including the “true” interaction intensities as latent variables without any restrictive parametric assumptions. Our simulation studies and real data analysis suggest the proposed approach achieves higher ranking accuracy, more interpretable identified peaks, and better consistency across biological replicates.

## 2 Model and Methods

### 2.1 Model setup

Most existing HiC peak calling methods recognize that distribution of the observed contact counts between a loci pair change with the distance between them. Following the setup in Duan *et al*. (2010); Ay *et al*. (2014), we group all genomic loci pairs from one chromosome based on their genomic distance. Let *x_im_* for *i* = 1,…,*n_m_,m* = 1,…*M* denote the observed Hi-C count for the *i*th pair of genomic loci with distance *d_m_* between each other. where *d_m_* increases with m. We are interested in *δ_im_*, an indicator variable denoting whether there is non-random physical contact between this loci pair. Using this indicator variable, our inference goal can be formulated as the multiple testing problem for *H_null,im_* : *δ_im_* = 0 *vs H_alt,im_* : *δ_im_* = 1 for *i* = 1,…, *n_m_, m* = 1,… *M*. For all peak detection methods in the literature, the null is rejected if the observed count is higher than expected under the null. But these procedures vary in terms of the parametric assumptions on the observed counts as described previously.

For *i* = 1,…, *n_m_, m* = 1,… *M*, we model the observed count *x_im_* with the following hierarchical model,

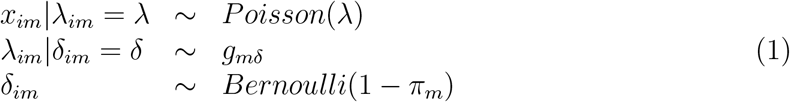

where *π_m_* is the probability for any loci pair with distance *d_m_* to be a non-peak, and *λ_im_* is a latent variable representing the “true” interaction intensity for this loci pair, whose distribution is *g*_*m*0_ under the null, and *g*_*m*1_ under the alternative. Marginally, the overall distribution of *x*im** can be written as the following two-group model

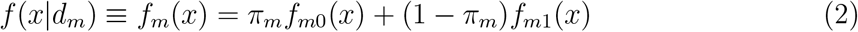

where *f*_*m*0_(*x*) and *f*_*m*1_(*x*) are the null and the non-null components, respectively.

We propose Empirical Bayes model of **Hi-C** (**EBHiC**), an empirical Bayes test for Hi-C peak identification with the following steps

1. Estimate the overall distributions *f*m** for *m* = 1,…, *M* jointly with spline-Poisson mixture models.
2. Estimate *f*_*m*0_ by approximating the ideal empirical null distribution based on *zero assumption.*
3. Rank the tests by local false discovery rate, and control the false discovery rate via thresholding.

The general empirical Bayes multiple testing framework was first proposed in Efron *et al.* (2001); Efrom (2004), and has become popular in genomic applications. In most of the previous literature on empirical Bayes test, the overall distribution of the test statistics are estimated using a nonparametric procedure such as Poisson regression over the histogram. The null distribution is typically estimated using a *central-matching* method based on the *zero assumption,* which essentially implies that the most of the tests are not significant. Zhang and Keles (2017) proposed to use spline mixture model to estimate the overall distribution of a continuous latent variable, and derive the empirical null using another natural consequence of the zero assumption. They then applied their proposed method on Binomial data arisen from the allele-specific ChIP-Seq binding detection. All the above works focused on one single two-group model. The current paper differs from the previous literature, because we study an ordered collection of two-group models indexed by *m* = 1,…, *M*. This is the our major methodological innovation. In what follows, we will present the elements of EBHiC in details.

### 2.2 Estimating *f_m_*(*x*) using smoothed EM algorithm

Similar to the observed count data *x_im_*, the latent variable *λ_im_* also follows a two-group model marginally for all loci pairs in group *m* ∈ {1,…,*M*},

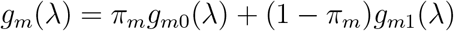

We propose to estimate *g_m_* using cubic B-splines. For a sufficiently large positive real number *U*, let *U*(*j* – *k* + 1)/*J* for *j* = 1,…, *J* + *k* +1 be equally spaced knots, and *B_j_*(*λ; k*) be the normalized k’th order B-spline defined on [*U*(*j* – *k* + 1)/*J*, *U*(*j* + 1)/*J*] such that 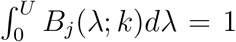. These basis are restricted within [0, *U*] and only the basis for *j* = 1,…, *J* + *k* – 3 are kept. Let *g_m_*(λ) be a smooth density function which we can approximate as:

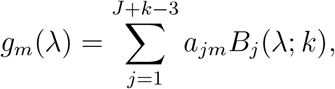

where *a_jm_* are non-negative coefficients summed to 1 for each of *m* = 1,…, *M*. So (2) is a spline-Poisson mixture model. It also follows that the marginal likelihood of the observed counts *x_im_* is given by

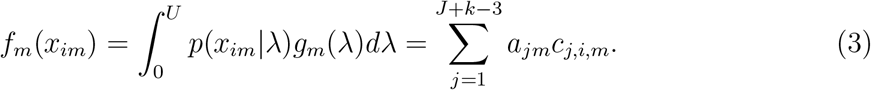

where 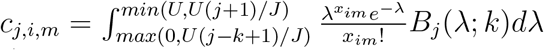 is a constant depending on no unknown parameters.

We use the same knots and B-spline basis for *m* = 1,…, *M*. The number of knots could be decided based on the data. Thus it is fixed for a given dataset. We then estimate the coefficients *a_jm_*’s by maximizing the likelihood of all observed data 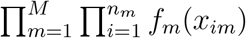 via a smooth Expectation-Maximization (EM) algorithm (Silverman *et al.*, 1990). The probability mass function in (3) for fixed m can be seen as a finite mixture distribution. Define *y_j,i,m_* as the indicator that *x_im_* is from component *j* ∈ {1,…,*J* + *k* – 3}, then *P*(*x_im_*|*y_j,i,m_* = 1) = *c_j,i,m_*, and *P*(*y_j,i,m_* = 1) = *a_jm_*. Since *f_m_* for *m* = 1,…, *M* is an ordered sequence of distribution, we also introduce a smoothing step using trigangular kernels to improve the smoothness of the estimated distributions, and facilitate information sharing across groups *m* = 1,…,*M* (See Supplementary Notes for more details on triangular kernels). To alleviate the issue with the different range of the counts across groups, we also add a power transformation on the latent variable λ_**im**_ (See Supplementary Notes for details).

The exact smoothed EM-algorithm is as follows.

1. *Initialization*. 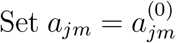 for *j* = 1,…, *J*+*k* – 3 and 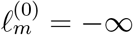 for *m* = 1,…,*M*.
2. *E-step.* For current coefficients 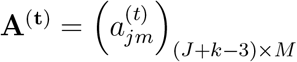,

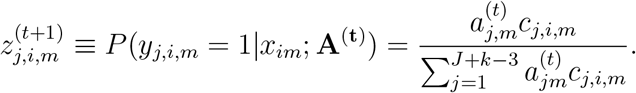
3. *M-step*. Raw estimate of the coefficients:

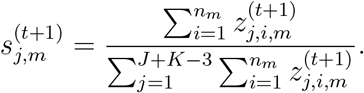

and let 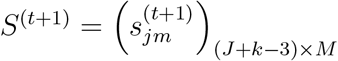
4. *S-step* Update the coefficient after smoothing:

a. Let 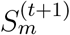 be the *m*’th column of 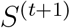, and 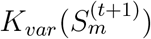 be a triangular kernel matrix defined on {1,…, *J* + *k* – 3} with fixed bandwidth and variable arms determined by 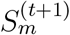, we first smooth the columns of 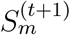 by

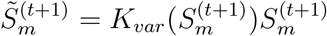
b. Let 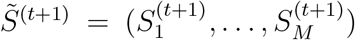, and *K_f_* be a fixed triangular kernel matrix defined on {1,…, *M*} with fixed bandwidth and arm. We update the coefficient matrix by

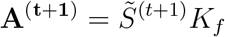
5. *Stopping rule.* Repeat steps 2-4 until the increase in the following estimated marginal likelihood is small in terms of both the absolute and the relative value for all *m* = 1,…,*M*.

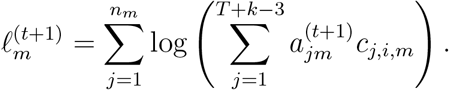 Stop when

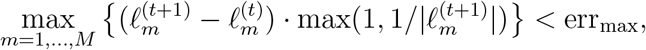

where err_max_ is a pre-specified control parameter.

### 2.3 Estimating *f*_*m*0_(*x*) and *π_m_* based on the zero assumption

For *m* = 1,…, *M*, we model *f*_*m*0_ as a Zero-Inflated-Negative-Binomial (ZINB) distribution with parameters (*p*_*m*0_,*μ*_*m*0_, *ϕ*_*m*0_) as its proportion of the extra zeros, and the mean and the over-dispersion parameter of the Negative-Binomial component. This is more flexible than the NB and ZINB models used in the literature as it allows different over-dispersion parameters across groups.

The observed count distribution for group m is the mixture of the null distribution *f*_*m*0_ and the non-null component *f*_*m*1_, respectively. It is often assumed that f_m1_ is zero in the bulk region of the overall distribution due to the identifiability issue. For z-scores, this bulk region is an interval around 0, and it is the set of non-negative integers below some threshold in our case. This is referred to as the *zero assumption* (Efron, 2012). Under this assumption and when the observed data are continuous and i.i.d., the null distribution can be estimated empirically using *central-matching* (Efron *et al.*, 2001). However, it cannot be directly applied to our discrete data.

Zhang and Keles (2017) observed another implication of the zero assumption. The ratio of the null and the overall distributions is close to a non-zero constant in the bulk region, and is close to zero if the non-null signals are detectable. In the case of a continuous latent variables, they proposed to estimate the empirical null by fitting a parametric density curve that minimizes the L2 norm of the derivative of ratio of the null and the overall densities. Following the same spirit, we propose to estimate the null models for our discrete data by minimizing the loss function

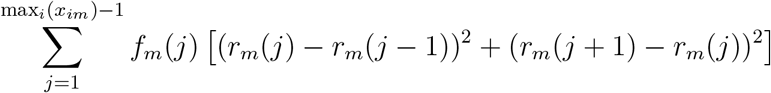

where *r_m_*(*x*) = *f*_*m*0_(*x*)/*f_m_*(*x*) is the likelihood ratio of the null and the overall model, and *f_*m*_* is estimated from our smooth EM algorithm. This is the discrete generalization of the curve fitting method in Zhang and Keles (2017), which replaces the derivative with the finite difference. The optimization is constrained with the assumption that the mean of the null model is smaller than the mean of the data in the same loci pair group. This is because our hypothesis testing problem is one-sided in the sense that the null is rejected only when the observed count is large enough. We remark that this formulation does not assert any additional assumptions other than the zero assumption commonly used in the literature of empirical Bayes modeling.

In theory, *π_m_* = 1/*max_x_*(*r_m_*(*x*)). In practice, we estimated it with 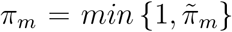 where 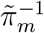 is the 0.98 quantile of the likelihood ratio for all observed counts in the null region. The null region is defined as the interval from 1 to the 0.9 quantile of the observed counts in group *m*, as it is unlikely to have more than 10% of the loci-pairs to be in meaningful physical contact.

### 2.4 Ranking the tests and controlling FDR using local false discovery rate

The local false discovery rate (locfdr) is defined as

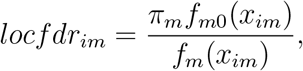

and has the interpretation of *P*(*H_null,im_* is true| data).

Let 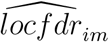 be the estimate of *locfdr_im_* by plugging-in the estimates of *f_m_*(*x_im_*), *f*_*m*0_ (*x_im_*) and *π_m_*. We reject the nulls for the loci pairs with small locfdr. Let 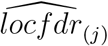 be the j’th (increasingly) sorted estimated locfdr value for all loci-pairs in all groups. Using the strategy used in (Zhao *et al.*, 2013; Zhang and Keles, 2017), FDR control can be achieved by

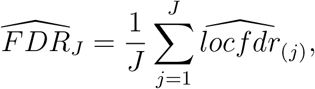

## 3 Results

### 3.1 Simulation Studies

We conduct a simulation study to examine the ranking accuracy of EBHiC. We simulate data from model (1) using the null parameter estimates from the analysis of the real data (Section 3.2) from chr1 of GM12878 with 10kb resolution after the following a few modifications. Without the loss of generality, we only used the null parameters for loci pairs with distance ≤ 1Mb to reduce the computational burden in simulation, and we simulated 2000 loci pairs for each distance group. We keep the null parameters (*μ*_*m*0_,*ϕ*_*m*0_,*p*_*m*0_) the same as the real data estimates, and multiply the proportions of the null *π_m_* by 0.95 to increase the number of true contacts. To investigate the effect of the strength of the true physical contacts on HiC peak calling, we model non-null distribution of the observed counts for each specific group m as a Uniform-Poisson mixture, where the expected values for the Poisson distributions follow *Unif* [1.1*μ*_*m*0_, (1.1 + *k*)*μ*_*m*0_]. Here *μ*_*m*0_ is the null parameter estimated from the real data, representing the mean of the negative binomial component of the ZINB null model, and *k* is a simulation parameter controlling the signal strength of the physical contact. Intuitively, the performance of any HiC peak detection algorithm should improve as k increases. We consider *k* = 0.25, 0.5,1, 2, and repeat the simulations for 20 replicates for each setting.

#### 3.1.1 EBHiC achieves higher ranking accuracy

We measured the ranking performance of each method using the area under precision-recall curve (AUPRC) as precision-recall curves are more appropriate for imbalanced data as ours where there is only a small proportion of loci pairs have physical contact. In Fig. 1, we find that EBHiC yields higher ranking accuracy than Fit-Hi-C in all simulation settings. When the signal strength increases, all methods perform better, and the gap between EBHiC and Fit-Hi-C decreases. It suggests that EBHiC is especially appropriate for situations where the signal is rare and weak.

**Figure 1:**
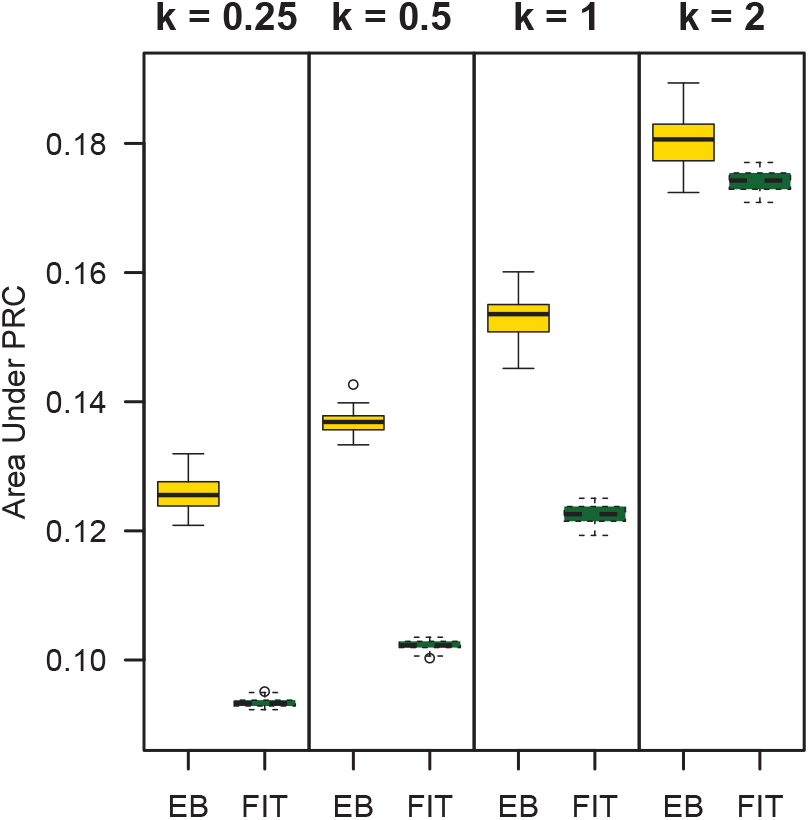
Simulation results: boxplot of area under the precision-recall curve for EBHiC (EB) and Fit-Hi-C (FIT)

#### 3.1.2 The power of EBHiC lays in the top candidate selection

While AUPRC is a widely accepted measure of the ranking accuracy of all candidates, we often only interested in a small proportion of the top candidates in the practice of HiC peak calling. In fact, previous works in the literature (Ay *et al.*, 2014; Schmitt *et al.*, 2016) seldom selected more than 1%-2% of the candidate loci pairs as the significant physical contacts. Thus we also investigate the accuracy of a given selected proportion of top candidates, prioritized by each method. We find that the top candidates ranked by EBHiC are more accurate than those by Fit-Hi-C, especially when only a small proportion of top candidates are concerned (Fig. 2). Even though Fit-Hi-C appears to catch up if more than 20% of the candidate loci-pairs are identified as significant physical contacts, selecting such high proportion of Hi-C peaks rarely happen in scientific practice.

**Figure 2:**
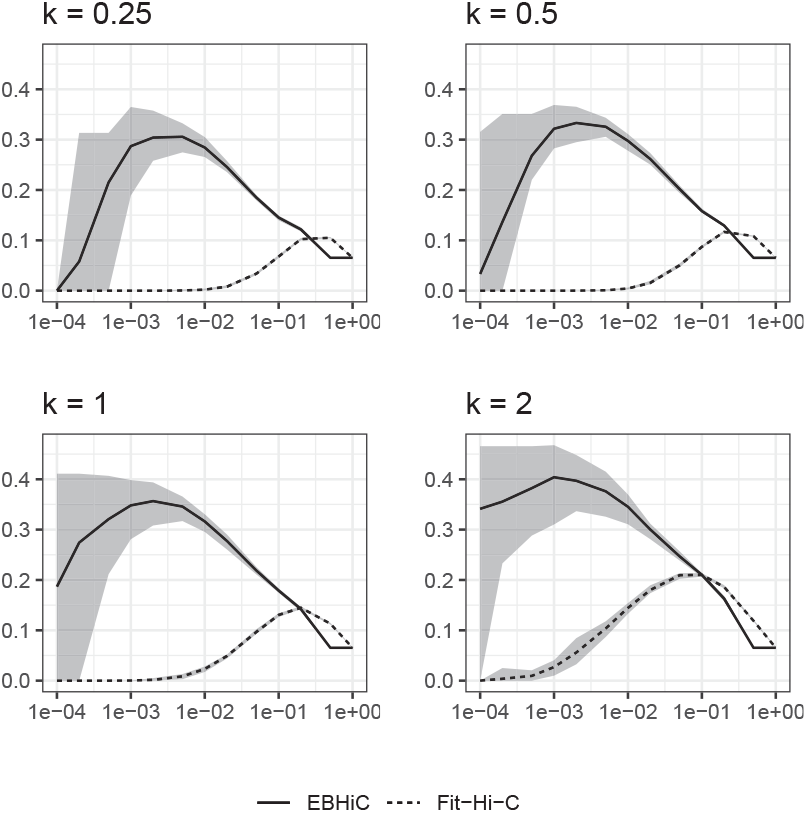
Simulation results: accuracy of selected top candidates (the ratio of the true positives and the number of selected) by EBHiC and Fit-Hi-C, respectively. The X-axis is the proportion of loci pairs selected in log scale. The Y-axis is the accuracy. The curves are the means, and the boundaries of the shaded areas are the 5% and 95% quantiles of the results from simulation replicates.

### 3.2 Real Data Analysis

In this section, we analyze the Hi-C data with 10kb resolution from IMR90 and GM12878 cells preprocessed as in Schmitt *et al*. (2016). Since most biological meaningful physical contacts at high resolution are between loci close to each other, we only focused on the intra-chromosome pairs with distance less than 2Mb, which is a common strategy taken in the literature of Hi-C peak detection (e.g., Schmitt *et al*. 2016).

### 3.2.1 EBHiC and Fit-Hi-C prioritize distinct types of loci pairs

For each dataset, we first examine the top selected loci-pairs by each method, and find that the overlaps between the top hits by EBHiC and Fit-Hi-C range between about 5% to 50%, and vary with the top proportion selected and cross the cells (Table 1). It suggests that EBHiC and Fit-Hi-C prioritize distinct types of pairs.

**Table 1:**
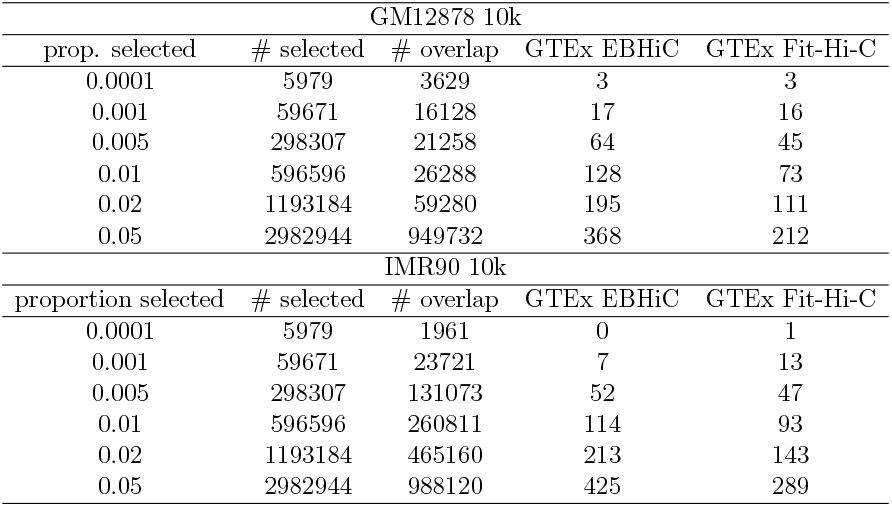
Comparison of the top ranked loci pairs by EBHiC and Fit-Hi-C. We select the same proportion of loci-pairs from each chromosome (prop. selected), and report the total number of selected pairs for the whole genome (# selected), the number of overlaps between EBHiC and Fit-Hi-C (# overlap), and the numbers of GTEx eQTL target-promoter pairs captured by EBHiC (GTEx EBHiC) and by Fit-Hi-C (GTEx Fit-Hi-C). Proportions larger than 0.05 are not considered as it is generally not expected to have more than 5% of the loci pairs are in biologically meaningful physical contact at high resolution such as 10kb in practice.

### 3.2.2 Higher proportions of EBHiC top loci-pairs overlap with GTEx eQTLs in relevant tissues

Next, we evaluate the accuracy of these top hits. Since there is no “ground truth” for long-range physical contacts of DNA, it is only possible to use relevant biological annotations from independent data type as benchmark. One potential “gold standard” of this kind is the database of the enhancers and the promoter of their target genes in relevant tissue. However, most of such databases are in fact computational predictions, some of which even use Hi-C data as input (e.g., Fishilevich *et al*. 2017). Experimentally validated enhancer-promoter pairs are rare. Another equally reasonable “gold standard” is the eQTL database for expressions in relevant tissues, as previous studies have shown that some trans-eQTLs are actually in physical contact with the promoters of the target genes in 3D space (Duggal *et al.*, 2013). We turn to GTEx database (Consortium *et al.*, 2017), and utilize the eQTLs from whole blood and EBV transformed lymphocytes to benchmark the analysis of GM12878, and those from lung and the transformed fibroblasts for IMR90. The p-value cutoff for eQTLs is 1 × 10^-8^. Overall, we find that EBHiC outputs hit more pairs of eQTL and target-promoter from GTEx than Fit-Hi-C in at most thresholds for both datasets.

### 3.2.3 EBHiC results are more consistent across replicates

We also examine the consistency in peak calling between biological replicates using the two pre-processed H1 cell replicate data with 40kb resolution from Schmitt *et al*. (2016). For a fixed proportion of top candidate loci-pairs from two biological replicates, we measure their consistency using Jaccard Index. In Fig. 3, we find that EBHiC leads to higher consistency cross biological replicates, especially when a small proportion of top candidates are identified as significant Hi-C peaks (e.g., < 1≤), which resonates with the findings in our simulation studies that the power of EBHiC lays in the improved ranking of the top candidates.

**Figure 3:**
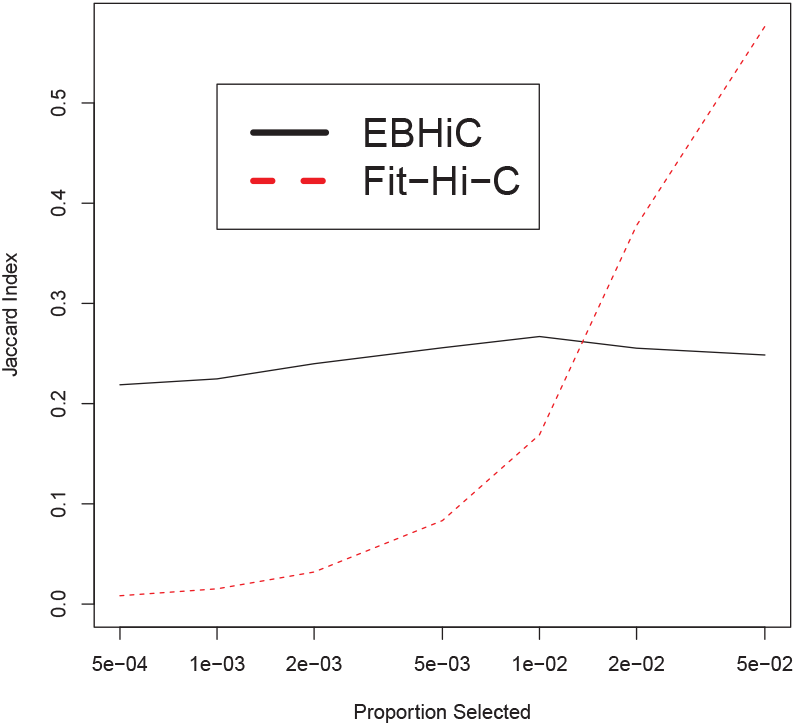
Real data analysis results: the Jaccard index of the selected top candidates from two H1 cell replicates, prioritized by EBHiC and Fit-Hi-C, respectively. The X-axis is the proportion of loci pairs selected.

## 4 Discussion

Despite the recent development in HiC analysis tools, the identification of chromatin interactions is still challenging due to the high dimensionality, sparsity and overdispersion entangled in the contact count matrix. Previous methods for HiC peak detection has not sufficiently addressed the over-dispersion of the count data by either imposing strong restrictive parametric assumptions or ignoring this issue completely.

In this paper, we propose EBHiC, an empirical Bayes model for HiC data modeling and peak identification. EBHiC estimates an overall probability distribution and a null distribution for the observed counts in each distance group, and identifies the peaks by ranking all loci pairs in all groups using their associated local fdr. The overall distribution of each group is modeled as a spline-Poisson mixture, and estimated jointly using a smoothed EM algorithm in which triangular kernels are applied enabling smooth probability distribution estimates and information sharing across nearby groups. The null distributions are estimated based on a natural consequence of the *zero assumption*. We conducted extensive simulation studies and real data analysis to evaluate the performance of EBHiC in terms of accuracy and consistency in peak calling.

EBHiC shares certain methodological similarity with Fit-Hi-C, but also possesses many critical differences. For Fit-Hi-C, its initial spline fit can be regarded as the smooth estimates of the expected values for the overall count distributions for each group, and the refit as the expected values for the null distributions. Then Fit-Hi-C proceeds to evaluating the statistical significance using p-values in a frequentist fashion. In contrast, EBHiC provides the smooth estimates of the whole probability distributions for each group under the overall and the null models, and evaluate the statistical significance using a principled empirical Bayes framework. Since the whole probability distributions are estimated, EBHiC also enables flexible data-driven estimates of the data over-dispersion, which cannot be done using the restrictive binomial models as in Fit-Hi-C.

EBHiC is a versatile HiC peak detection tool that is agnostic on the HiC data preprocessing pipeline, as long as the contact counts are provided, which does come with the price of losing the opportunity of accounting for factors such as mappability, GC content, and multi-mapping reads in the peak calling step. However, these problems could be alleviated by incorporating bias correction procedures such as ICE (Imakaev *et al.*, 2012) and multi-read mapping tools such as mHi-C (Zheng *et al.*, 2018) in the preprocessing step.

Similar to the majority of the existing HiC peak identification tools, EBHiC does not take into advantage of the spatial information in the contact matrix. One potential extension of EBHiC along this direction is to replace the bottom Bernoulli layer of model (1) with a binary Markov Random Field (MRF) as in Xu *et al*. (2015). Instead of implementing a fully Bayesian estimate for this hidden MRF, we could potentially take advantage of the Bayesian interpretation of locfdr to execute a faster mean field approximation. Another possible extension of EBHiC by incorporating the spatial information lays the connection between the inference goals of HiC peak detection and TAD identification, which are finding the significant physical contact between individual loci pairs, and the large scale regions enriched with true physical contact. Our proposed empirical Bayes framework enables the possibility of the joint peak detection and TAD identification by adding additional layers of Bernoulli latent variables in (1) to model the TAD structure, and the loci pairs within or between TADs.

## Acknowledgements

We thank the Holland Computing Center (HCC) at UNL for computation resources and technical supports.

## Funding

QZ and YL’s research has been supported by NSF ABI (Award# DBI-1564621), NSF EPSCoR (RII) Track II (Award# OIA-1736192) and NU Collaborative System Science Seed Grant to QZ. ZX’s research was partially supported by Layman Seed Award from UNL.

## Supplementary Notes

### Implementation details

**Power transformation of** λ

To borrow strength across the bin pair groups indexed by *m* = 1,…, *M*, the knots and thus the spline basis for *m* = 1,…, *M* are the same. In practice, we set

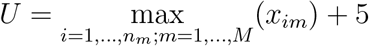

In HiC data, however, the maximum of 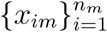 may decrease with *d_m_* dramatically, which cause the effective numbers of basis,i.e., the number of basis that covers [0, max_*i*=1,…n_m__ (*X_im_*)], for larger *m* may be much smaller than *J* + *k* – 3. To alleviate this problem, we actually use the splines to model the distribution of λ^*t*^ where *t* ∈ (0,1]. Then the overall density *g_m_*(λ) becomes

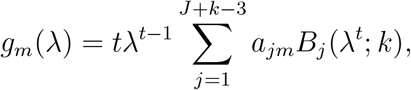

and the constants in *c_j,i,m_* are now

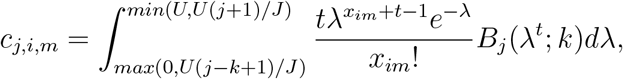

The power for the transformation, *t*, is selected based on the data using the following rationale.

Let *x_max,m_* = max_*i*=1,*n*_*m*__ (*x_im_*), then *U* = max(*x_max,m_*) + 5. Since the equally spaced knots on [0, *U*] are used, the proportion of the effective spline basis for a case *m* = *m*_0_ is roughly *x*_*max,m*_0__/ max(*x_max,m_*). Motivated by this, it is desirable to have the proportion of the effective basis to be larger than a threshold (e.g., 0.2) in the most extreme case after the power transformation, i.e.,

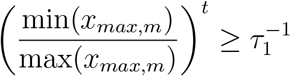

which implies

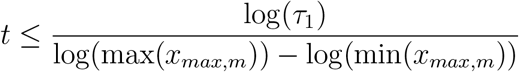

On the other hand, if *t* is too small, a large proportion of the basis will fall between [0,1],and become less useful. Thus we suggest choose *t* such that

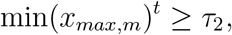

which implies

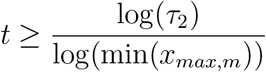

For prespecified parameters *τ*_1_,*τ*_2_ > 0, we choose *t* as the average of the above upper and the lower bounds of *t*. Even when there is no *t* that satisfies both conditions, the average is still a balanced choice. In practice, we let (*τ*_1_,*τ*_2_) = (3, 4).

### Triangular Kernels

For arm length a and bandwidth *h*, a discrete triangular kernel (Kokonendji *et al.*, 2007) on {1,…, *N*} is defined as

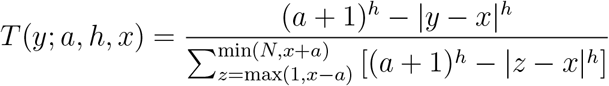

for |*y* – *x*| ≤ *a* and *x, y* ∈ {1,…, *N*}.

For the fixed (*a, h*), a triangular kernel matrix is defined as

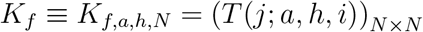

Its column sums are 1. We use this fixed triangular kernel matrix to smooth the rows of 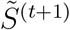 to enhance the similarity of *f_m_*’s for adjacent groups.

For the smoothing the columns of *S*^(*t*+1)^ in the smoothed EM algorithm, we adopt the following triangular kernel with variable arm length. In details, let *b* ∈ (0,1) and *w* = (*w*_1_,…, *w_N_*)^*T*^ be an estimated discrete distribution on {1,…, *N*}. We define the variable arm triangular kernel matrix as

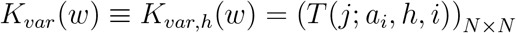

where

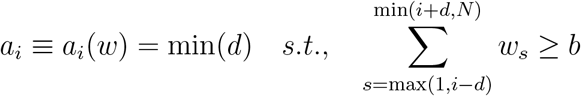

This kernel allows more aggressive information sharing and smoothing of the probabilities at where the data is scarce, and does the opposite in the bulk regions. In the literature, nonparametric density estimation with varying smoothness is usually accomplished by a kernel density estimator with adaptive or variable bandwidth (Givens and Hoeting, 2012). Our application is different, as we are smoothing the current estimate of the distribution instead of smoothing the raw data directly. We find that manipulating the arm length parameter a is more intuitive as it is closely related to the window length of moving window based smoothing.

